# CDR3 and V- genes show distinct reconstitution patterns in T-cell repertoire post allogeneic bone marrow transplantation

**DOI:** 10.1101/2020.03.05.977900

**Authors:** N Tickotsky-Moskovitz, S Dvorkin, A Rotkopf, A.A. Kuperman, Sol Efroni, Y Louzoun

## Abstract

**Background:** Restoration of T-cell repertoire diversity after allogeneic bone marrow transplantation (allo-BMT) is crucial for immune recovery. T-cell diversity is produced by rearrangements of germline gene segments (V (D) and J) of the T-cell receptor (TCR) α and β chains, and selection induced by binding of TCRs to MHC-peptide complexes. This diversity can be measured by many measures. We here focus on the V gene usage and the CDR3 sequences of the beta chain. We compared multiple T-cell repertoires to follow T cell repertoire changes post allo-BMT in HLA-matched related donor and recipient pairs.

**Results:** Our analyses of the differences between donor and recipient complementarity determining region 3 (CDR3) beta composition and V-gene profile show that the CDR sequence composition does not change during restoration, implying its dependence on the HLA typing. In contrast, V gene usage followed a time-dependent pattern, got following the donor profile post transplant and then shifting back to the recipients’ profile. The final long-term repertoire was more similar to that of the recipient’s original one than the donor’s.; some recipients converged within months while others took multiple years.

**Conclusion:** Based on the results of our analyses, we propose that donor-recipient V-gene distribution differences may serve as clinical biomarkers for monitoring immune recovery.

## Background

Over the last decades, bone marrow transplantation is being used to treat numerous malignant and non-malignant diseases in which a renovation of hematopoietic stem cells is needed (1–3). Immune reconstitution and regain of function after allogeneic bone marrow transplantation (allo-BMT) relies primarily on the homeostatic proliferation of mature T lymphocytes (T-cells) and is slow and complex (1,4–6). T-cell reconstitution can be influenced by several factors such as patients’ age, human leukocyte antigen (HLA) mismatches, source of stem cells, and the occurrence of graft versus recipient disease (GVHD)(5,7–9).

Complete reconstitution of the T cell compartment after allo-BMT occurs through two distinct pathways: an initial peripheral expansion and a later T cell production by the thymus (10). In the early post-transplant period (soon after transplant,day+30 or later), adoptively transferred donor T cells or recipient T cells that survive conditioning expand in the periphery (8). In the later post-transplant period naive T cells are produced *de novo* in the recipient’s thymus (11), which is seeded by lymphoid progenitors derived directly from the graft or arising from donor hematopoietic stem cells (10).

T-cell reconstitution can be tracked by the T-cell receptor (TCR) repertoire diversity. This diversity is driven by rearrangements of the TCR α and β chains gene segments (V (D) and J), formation of the junctions between them (12), and the following selection through binding to MHC-peptide complexes. A large part of repertoire diversity and its binding to target MHC-peptide are determined by the beta chain (TCRβ) (13). During segment joining, nucleotides are inserted and deleted at the junctions between pairs of rearranging genes (12), forming the highly variable complementarity determining region 3 (CDR3) that is directly implicated in antigen recognition (12,14–16). Given its importance in antigen binding, the CDR3 beta sequence is often used as a measure of repertoire diversity (17,18).

A parallel measure of repertoire diversity is V gene usage. The V gene usage is affected by the genetic composition of the V locus haplotypes, both in gene usage and in gene expression levels (19,20). While there are consistent patterns of V gene usage in rearranged TCRs, with some V genes consistently expressed at a much higher level than others (16), almost every sample donor has unique V haplotypes, with differences in allele composition (21,22). Thus, aimed to find out if V beta usage can be treated as a marker for the origin of the T cells.

To measure CDR3 beta sequence composition and V gene usage, we studied the repertoire of donor-recipient pairs at different time points. Repertoire next-generation immune-sequencing (often denoted rep-seq) (23) supplies relatively detailed information on T cell repertoires (22,24–27) and has been applied to tracking T-cell reconstitution after allo-BMT (2,3,7,28–30). Current analytical approaches for the comparative analysis of TCR repertoires are mainly based on statistical inference and include repertoire diversity, repertoire overlap, V and J-gene segments usage similarity, and amino acid composition of the CDR3 (31,32). A recent quantitative work on the organization of post allo-BMT repertoire compared donor-recipient TCRβ V-J repertoires by self-similarity mathematical measures and found differences between recipients who had GVHD and those who did not (33).

## Results

We characterized each repertoire by its CDR3 beta or V-gene distribution and then compared these characteristics between all repertoires (See flowchart of analyses in Figure 1). Our results show that V gene usage and CDR3 sequences composition reflect the integration process of the transplanted cells in the recipient’s immune system. Our findings suggest that the regenerating repertoire is influenced at different time points by both donor cells and recipient cells and that this dual contribution results in convergence towards the original homeostatic distributions.

**Figure 1.**
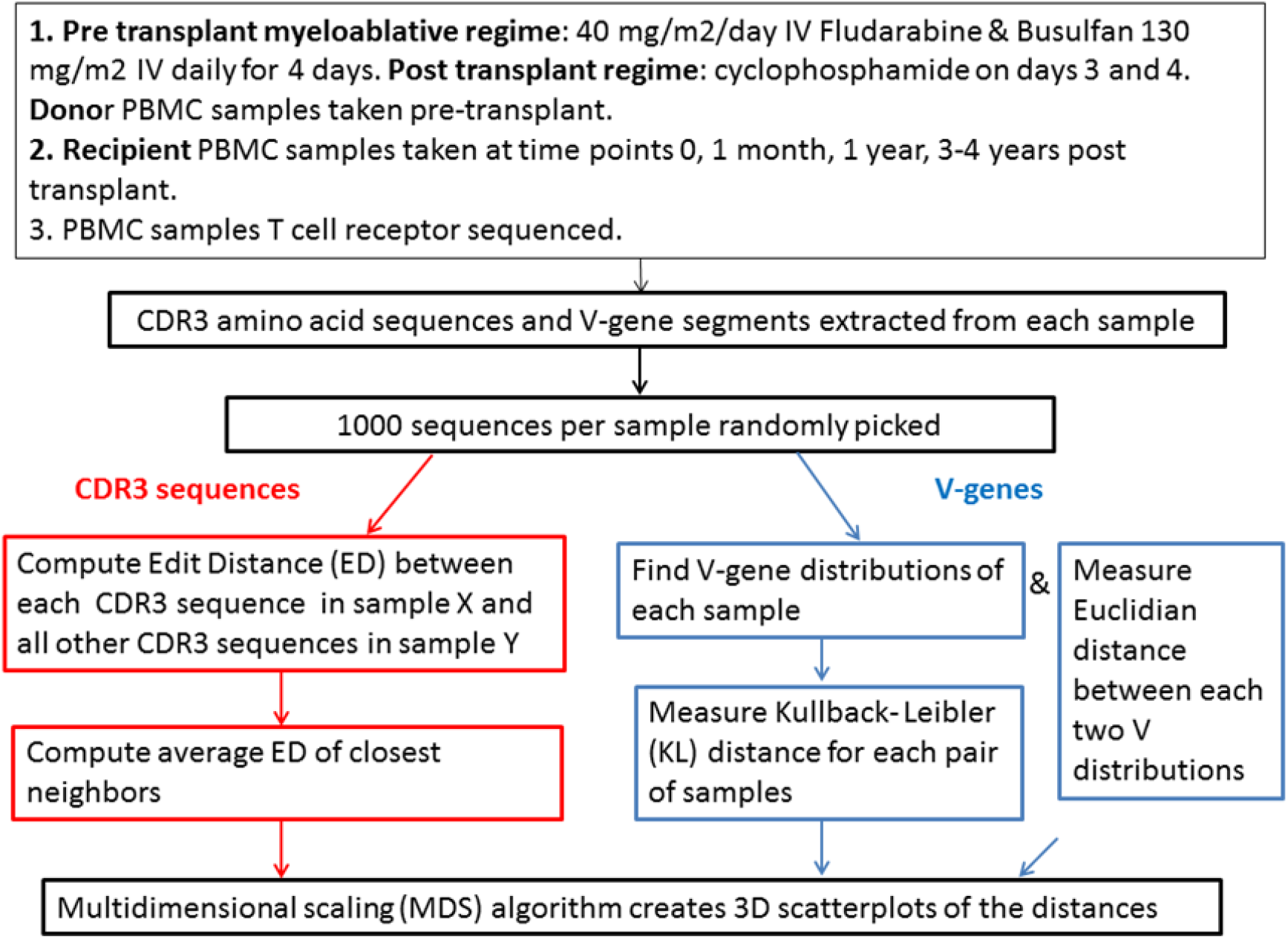
Flowchart of data analysis. TCR repertoires are sampled to a constant number of clones, and the samples are analyzed through their CDR3 beta receptor sequences and V gene distribution. We study each aspect by itself, and show a differential development of the two.

### V usages among samples

V usage is affected by the host genetics, but also by the conditioning of the patients. We measured the distance between pre-transplant (and pre-ablation) recipient repertoires, post-transplant recipient repertoires at multiple time points, and the donor repertoire. To avoid sampling biases, the same number of clones in each sample was used. We have estimated the distribution with a 1,000 clone sample (See Figure S5 for validation that this sample is large enough).

The diversity of the repertoire is often defined using clone size distributions and estimates on the repertoire diversity based on such distributions (26,34). Although clone size distributions help in estimating the immune potential of the regenerating repertoire, such measures are sensitive to sampling depth. Furthermore, the clone size distribution does not measure the difference between clones. To address that, we define the “diversity” of samples as the average distance of each clone to the most similar clone in the same sample. A short distance implies clones are close to one other, while a large distance implies that clones are uniformly distributed and not in close groups.

The similarity between each pair of samples, i.e., the distance between them,was computed. To account for the bias in V-gene usage over the entire repertoire (explained in the methods section), the symmetrized Kullback Leibler (KL) (35) divergence was used. In contrast with the Euclidean distance, the KL divergence is sensitive to changes in rare V genes.

*Figure 2*. A. V-

The distances were clustered using an average link hierarchical clustering. The samples show a clear division into two groups, each showing intra-group similarity and distinction from the other group, as demonstrated in the dendrogram and distance heatmap (Fig. 2A and 2B) and in the distance distribution (Two groups of distances in each color Figure 2C) (Student’s t-test of distances within the two main clusters and between clusters, p-value< 1.e-10). The similarity within a group of samples of the same time point was much lower than the similarity within recipients (Fig. 2A).

**Figure 2.**
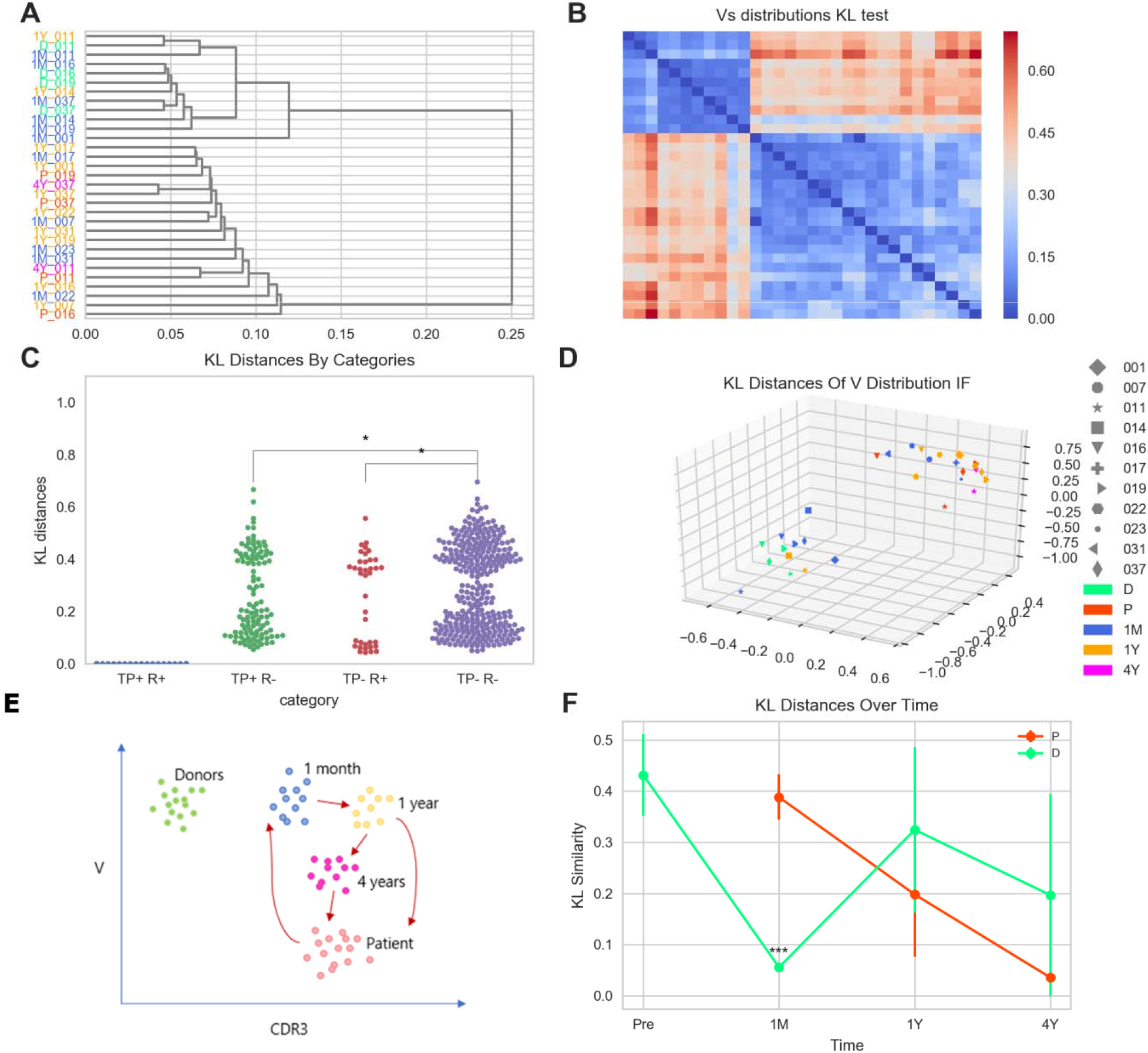
**A.** V-gene hierarchical clustering. KL distances between samples were clustered with ‘single’ method. The first half of the labels indicates one of the five states: Donors (D), Pre-transplant (P), 1-month post (1M), 1-year post (1Y) and 3-4 years post-transplant (4Y). The second half of the labels indicates the ID of the recipient. Labels are colored according to time points (TP). **B.** KL distance calculations were tested on uniform samples of 1000. In the KL V-gene distribution heatmap each cube represents the KL result from two distributions of V-gene usage. Blue indicates smaller KL values, i.e., more similar samples. Samples are positioned by the order of the dendrogram leaves from 2A. **C.** A swarm plot (i.e., a categorical scatterplot) of KL distances from 2B, grouped into 4 categories: ‘same TP, same recipient’ (time point= TP) (Blue), ‘same TP, different recipient’ (Green), ‘different TP, same recipient’ (Red) and ‘different TP, different recipient’ (Purple). Each point represents a sample. The Student’s t-test’s p-value is indicated by * (p-value < 0.05).**D.** A multi-dimensional scaling analysis projected the samples into a three-dimensional space, (maintaining the distances as in Fig 2B), with each point representing a sample. The analysis shows that the groups are defined by time point and not by recipient (Figure 2D). Different time points are shown in different colors and different sample IDs are indicated by a variety of shapes. We can see that each group contains different individuals from the same time point. **E.** A graphic illustration of the repertoires’ reconstitution: One-month post transplantation, some recipients’ repertoires share similarities with the donors and some are more similar to their own pre-transplant repertoires. After only one year, most repertoires from the slower-rate group become more similar to before-transplant repertoires. Then, in the 3-4-year group all recipients are closer to their pre-transplant repertoires than to their donor’s repertoire. **F.** KL distances of four recipients (011, 037, 016, 019) from their original pre-transplant repertoires and from their donors’ repertoires as a function of time. Recipients’ curves are in red, donors’ are in green.

A multi-dimensional scaling (MDS) analysis illustrates sample clusters in a three-dimensional space (maintaining the distances as in Fig 2B). Each group contains different shapes (each shape depicts one individual) of the same color (each time point has a distinct color). The groups are defined by time point and not by the recipient (Fig. 2D). The difference between donors in general and pre-transplant patients may represent either the effect of conditioning or the disease.

As shown in Figure 2F, donors and pre-transplant recipients are the most distant (i.e. non-similar). One-month post-transplant the repertoire of most samples and all CMV-negative recipients but one is very similar to the donors and different from the pre-transplant (Table2).

In all other samples, two distinct groups can be observed, which are both equally distant from the donor and the recipient, but far from the first group of one-month post-transplant. However, all groups slowly converge after one to four years towards a new distinct repertoire which is more similar to the original recipient’s than to the donor’s (Fig. 2F, p-value< 0.05 for all pairwise time point comparisons and for ANOVA on all time points). The results are consistent for each patient separately (Fig S6). We did not observe any effect of GVHD on the evolution of V-gene repertoire (Table 2). We performed a similar analysis using the regular Euclidean distance between repertoires, which yielded similar but less distinctive results, suggesting that the KL distance is indeed a better measure for V-gene similarity (Figure S7 A and B).

**Table 1.**
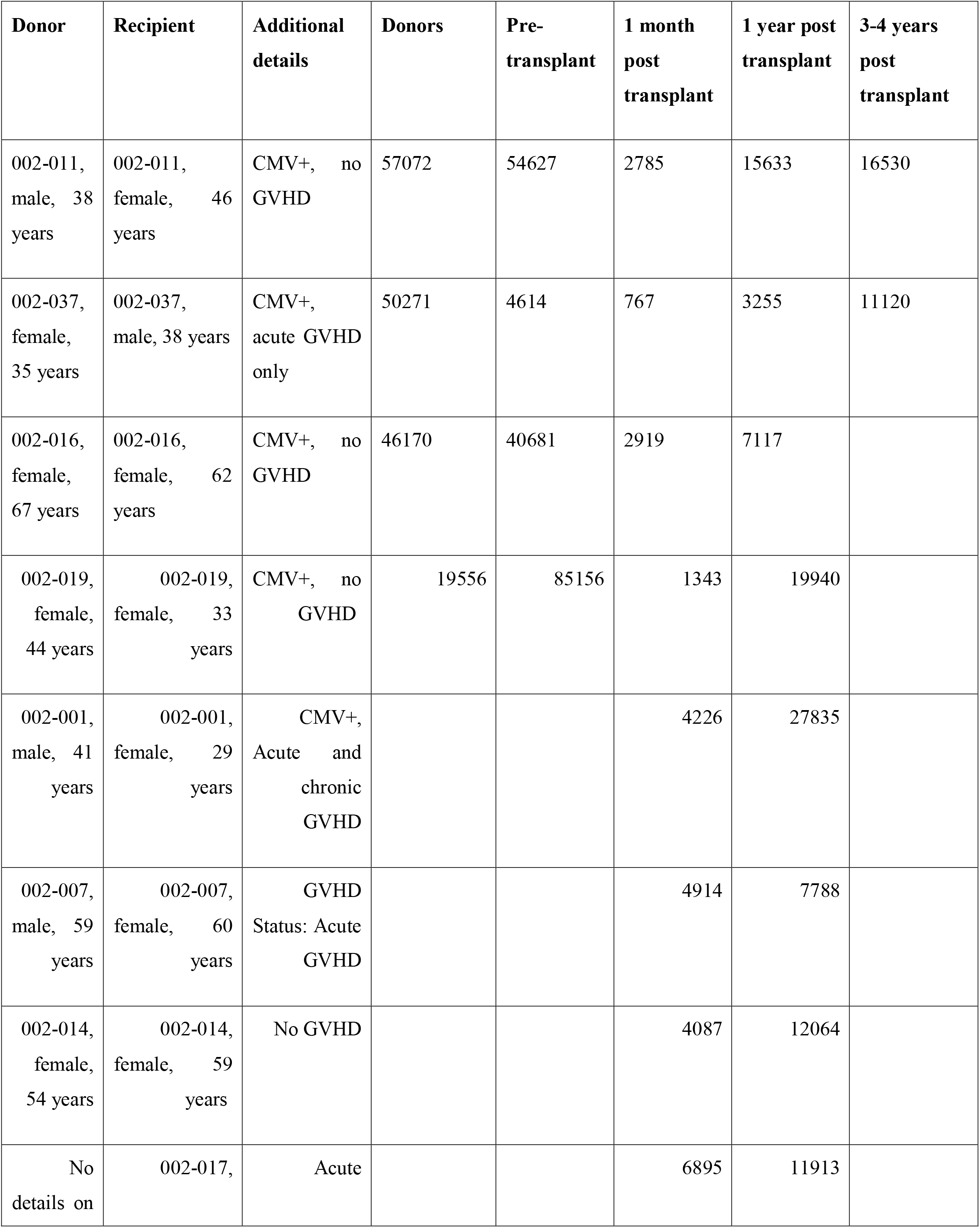
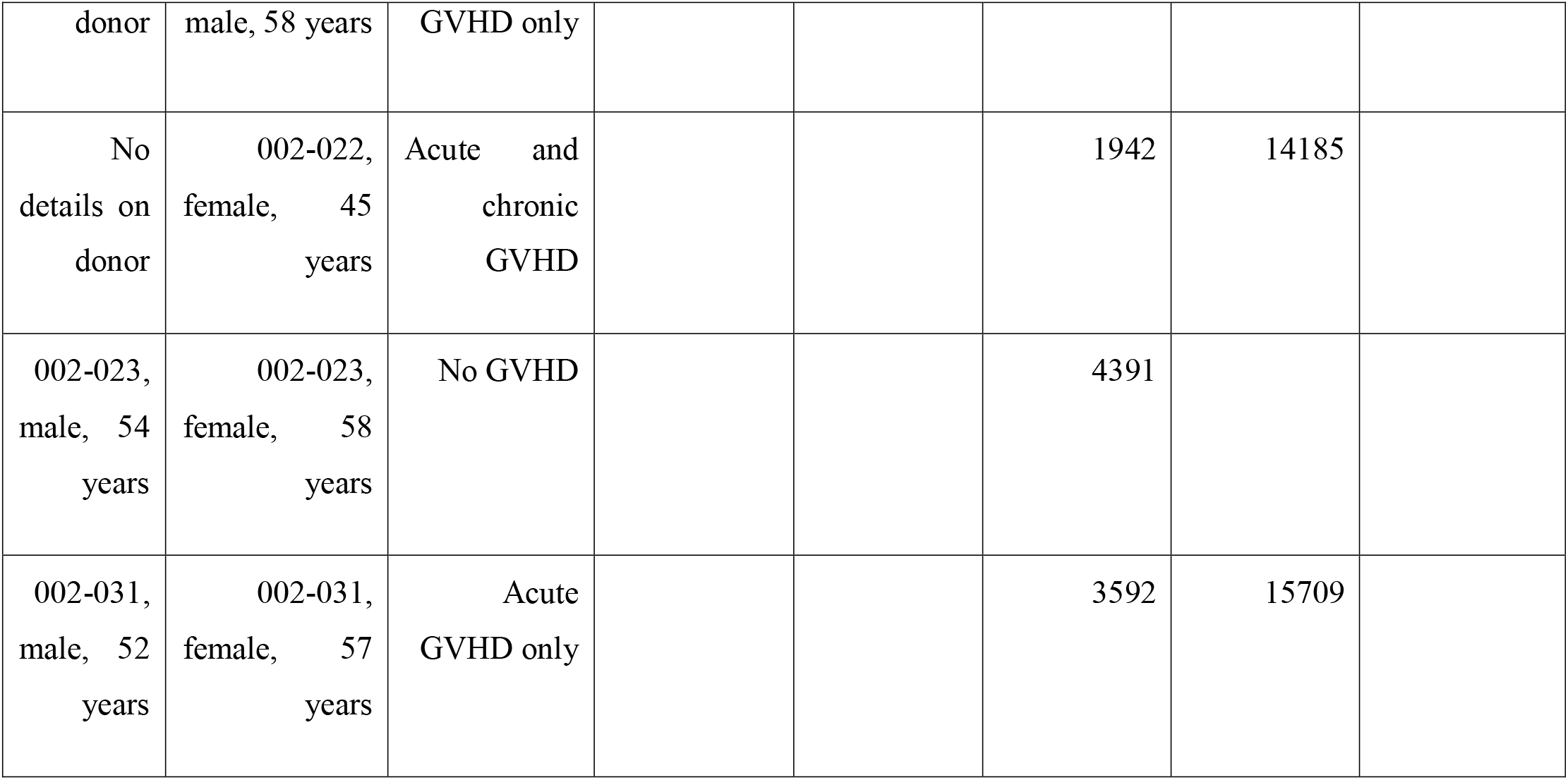
Donor-recipient pairs. The table lists each recipient with his/her donor. For seven recipients no donor repertoire was available. The numbers in the time point columns show the number of sequences in each file.

| **Donor** | **Recipient** | **Additional details** | **Donors** | **Pre-transplant** | **1 month post transplant** | **1 year post transplant** | **3-4 years post transplant** |
| --- | --- | --- | --- | --- | --- | --- | --- |
| 002-011, male, 38 years | 002-011, female, 46 years | CMV+, no GVHD | 57072 | 54627 | 2785 | 15633 | 16530 |
| 002-037, female, 35 years | 002-037, male, 38 years | CMV+, acute GVHD only | 50271 | 4614 | 767 | 3255 | 11120 |
| 002-016, female, 67 years | 002-016, female, 62 years | CMV+, no GVHD | 46170 | 40681 | 2919 | 7117 |  |
| 002-019, female, 44 years | 002-019, female, 33 years | CMV+, no GVHD | 19556 | 85156 | 1343 | 19940 |  |
| 002-001, male, 41 years | 002-001, female, 29 years | CMV+, Acute and chronic GVHD |  |  | 4226 | 27835 |  |
| 002-007, male, 59 years | 002-007, female, 60 years | GVHD Status: Acute GVHD |  |  | 4914 | 7788 |  |
| 002-014, female, 54 years | 002-014, female, 59 years | No GVHD |  |  | 4087 | 12064 |  |
| No details on | 002-017, | Acute |  |  | 6895 | 11913 |  |

|  |  |  |  |  |  |  |
| --- | --- | --- | --- | --- | --- | --- |
| donor | male, 58 years | GVHD only |  |  |  |  |
| No details on donor | 002-022, female, 45 years | Acute and chronic GVHD |  |  | 1942 | 14185 |
| 002-023, male, 54 years | 002-023, female, 58 years | No GVHD |  |  | 4391 |  |
| 002-031, male, 52 years | 002-031, female, 57 years | Acute GVHD only |  |  | 3592 | 15709 |

**Table 2.** Description of recipient’s samples characteristics.

| ID | GVHD | CMV | Time post- transplant |
| --- | --- | --- | --- |
| 011 | X | V | 4Y |
| 037 | Acute | V | 1Y |
| 016 | X | V | 1Y |
| 019 | X | V | 1Y |
| 001 | Acute+ chronic | V | 1Y |
| 014 | X | X | - |
| 007 | Acute | X | 1M |
| 023 | X | X | 1M |
| 031 | Acute | X | 1M |
| 017 | Acute | x | 1M |
| 022 | Acute+ chronic | X | 1M |

#### CDR3 composition among samples

The CDR3 beta distance distribution follows the recipients’ repertoire much more closely than the V-gene distribution Specifically, the similarity between the CDR3 sequences of samples from the same recipient is higher than between CDR3 sequences from different recipients at the same time point (both are more similar than CDR3 sequences from different recipients and different time points) (Student’s t-test p-value < 0.001), as can be seen in Figure 3A and 3B. Moreover, the repertoires of each donor and the appropriate pre-ablation recipient are more similar to each other than those of different donors, suggesting that the CDR3 sequence composition is indeed determined by HLA (either directly or through the peptide composition), since donor and recipients are fully matched.

**Figure 3.**
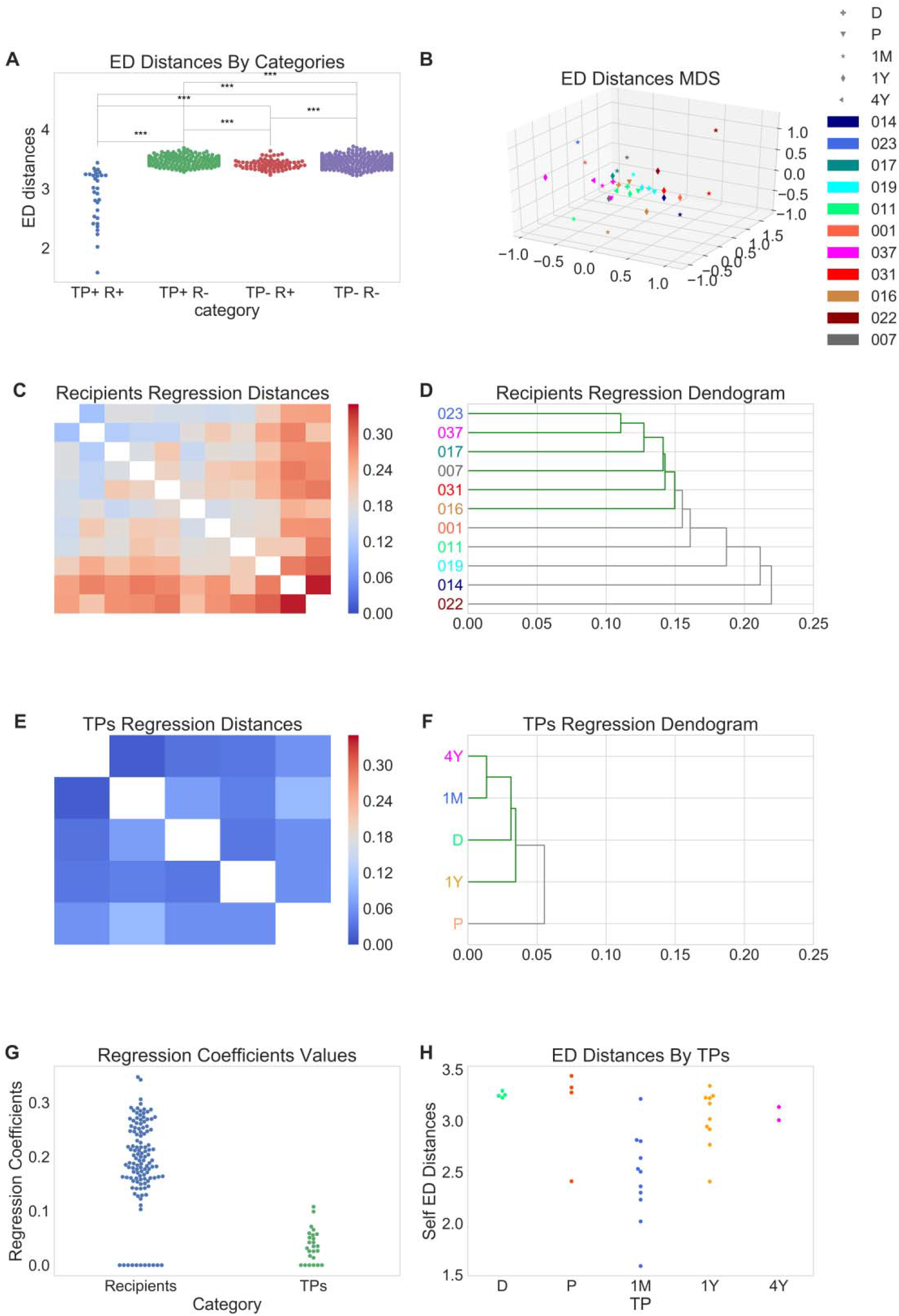
The main feature of the developing CDR3 repertoire is not a shift from the donor to the recipient (since there was no large difference to start with), but rather a change in CDR3 sequence composition (i.e., distances between clones). **A**. CDR3 Edit Distance by categories. Swarm plot of ED distances from 2B, grouped into 4 categories: ‘same time point’ (time point = TP), ‘same recipient’ (Blue), ‘same TP, different recipient’ (Green), ‘different TP, same recipient’ (Red) and ‘different TP, different recipient’ (Purple). **B.** MDS of the ED distance matrix; each point represents a sample. Different IDs are shown in different colors and different timeline states are indicated with five shapes. **C.** Regression values heatmap of all recipients’ distances. Recipients are ordered by Figure 3D leaves. **D.** Dendrogram of the values from Figure 3C. Label colors match those of the IDs from Figure 3B. **E.** Regression values heatmap of all time-points distances. Time points are ordered by Figure 3F leaves. **F.** Dendrogram of the values from Figure 3E. Label colors match those of time points from Figure 2D. **G.** Swarm plot shows the regression distances from C and E of the recipients and time points. Each point in the swarm plot represents a sample. When comparing the time points, repertoires at all post transplants time points are similar to each other, while both donor and recipient pre-transplants repertoires are farther away, as illustrated by the density of the post-transplant clusters as opposed to the dispersity of the pre-transplant cluster. **H.** Swarm plot of self ED distances by time points, colors match those of time points from F. Each point in the swarm plot represents a sample.

To test whether the distance in CDR3 composition was derived from the difference between recipients or from the reconstitution process, we separated the distance between samples (See Methods) into positive components based on the differences between recipients and components determined by the time points. The distance between recipients is larger than the distance between the time points (Figure 3C-3G). When comparing recipients, most have similar distances between each other, except for recipient 022 (Figure 3C, 3D), who shows a unique combination of positive CMV status, acute and chronic GVHD and a very low TCR count. In summary, repertoires at all posttransplant time points are similar to each other, while both donor and recipient pre-transplant repertoires are farther away (as shown in Figure 3E and illustrated by the density of the post-transplant clusters as opposed to the dispersity of the pre-transplant cluster in Figure 3F).

Using the self-distance as a diversity measure, the main feature of the developing CDR3 repertoire is not a shift from the donor to the recipient (since there was no large difference to start with), but rather a change in the CDR3 diversity (Figure 3H), which drastically decreases following transplant, and then converges back to the pre-transplant level. This self distance can be used in combination with the V gene usage distance as a clear measure of the repertoire reconstitution.

## Discussion

We propose a novel dual method for the analysis of TCR reconstition following tranplants. Each post-allo-BMT TCRβ repertoire is characterized by its CDR3 beta sequences and Vbeta-gene distribution. Then a distance is defined on the CDR3 sequences and V gene usage between samples. We have showed that V-gene and CDR3 sequences compositions, when tracked separately, can provide information on the reconstitution of the TCR repertoire.

V genes followed a time-dependent pattern; following transplant they shifted from the recipient to the donor profile and then back to the recipient’s profile. This is in accordance with Kanakry et al.(1) CDR3 beta sequences, on the other hand, remained constant within a recipient or its donor, suggesting an HLA-driven difference between hosts, with the main change being the decrease in diversity following transplant and a later expansion back to original levels.

The above described reconstitution patterns may reflect the two pathways of T-cell regeneration that act at different time points post-transplantation: The initial peripheral thymic-independent pathway (5,7,8) expands mostly the CD8+ memory T cells from the mature donor graft, which have previously encountered pathogens and are less dependent upon recognition of self MHC–peptide complexes for survival (5–8). In our analysis, CDR3 diversity decreased at this stage, in accordance with the shrinkage and skewing of the TCR repertoire in the initial wave of recovery leads to (10).

Interestingly, a dominance of pre-transplant CDR3 beta sequences was apparent. This finding is confusing, since, given that recipients had myeloablative conditioning, it is unlikely that residual recipient T cells had survived. This dominance of pre-transplant CDR3 beta sequences may result from recipient-derived selection applied in the course of peripheral expansion in response to viral antigens, with cytomegalovirus (CMV) being a known driving force in V usage for CMV+ patients (36), whereas non-CMV positive recipients likely respond to other viral antigens which may be recognized by similar T cells in the donor that would have existed in the recipient. The duration of this transitional pathway varies significantly among recipients.

In the second stage, which may take between less than a year and a few years, a thymic-dependent pathway is activated in the majority of patients(5,7), in which naïve T cells from progenitors of donor origin are increasingly produced *de novo* in the recipient thymus (5,7,8,37,38). Recovery through the thymic-dependent pathway accounts for the durable and clonally diverse reconstitution of the T-cell compartment (8,10), and has been found also in older patients (8). The timing of this process probably accounts for our observation that one-year post-transplant, V-gene segments in most samples and all CMV-negative recipients but one formed two distinct groups which were both equally distant from the donor and the recipient, but far from the group of one-month post-transplant.

V-gene composition differed significantly between pre-transplant recipients and their donors, probably because both disease conditions and treatments had affected the recipients’ repertoires. We suggest that the shift of V-genes from patient to donor profile one-year post-transplant reflects the thymic *de novo* production of naïve T cells from *donor origin* in the six to twelfth months post-transplantation. This has also been suggested by Meier et al.(2019) (33).

Timely reconstitution and regain of function of a donor-derived immune system is of utmost importance for the recovery and long-term survival of patients after allogeneic hematopoietic stem cell transplantation. In fact, the slow T cell reconstitution (6,8) is regarded as the primary cause for deleterious infections, the occurrence of graft-versus-recipient disease, and relapse (9,39). It takes weeks to months to produce naive T cells from infused stem cells and a plateau level of thymic output is reached only after at one to two years post-transplantation (11), rebuilding only completed after two to three years (11). We show that the transfer back to a repertoire similar to the pre-transplant one has different rates: in some patients, the convergence back is fast while in others it is quite slow. Donors and pre-transplant recipients’ repertoires were the most distant. This is in accordance with the findings of Buhler et al.(40), who studied a large cohort of 116 full chimeric recipients and their respective HSC donors pre-transplant and at one-year post-transplant. The recipients’ repertoire one-year post-transplant was very different from the donors’, with only a few overlaps. This finding led them to suggest that a new repertoire can be reconstituted at any age through thymic dependent or independent pathways (40).

As has been previously shown (33), the final long-term repertoire was more similar to that of the patient’s original one than the donor’s. We suggest a graphic illustration of the repertoires’ reconstitution that is based on the results of our analyses, as shown in Figure 2E: One-month post-transplantation, some recipients’ repertoires share similarities with the donors and some are more similar to their own pre-transplant repertoires. After only one year, most repertoires from the slower-rate group become more similar to before-transplant repertoires. Then, in the 3-4-year group, all recipients are closer to their pre-transplant repertoires than to their donor’s repertoire.

With recent advances in hematopoietic cell transplants, immune-monitoring is of major importance to identify patients at risk and provide them with early immune interventions (9). For instance, patients with low TCR diversity are at risk for hematopoietic cell transplantation-related complications such as GVHD and could benefit from adjuvant immunotherapies, such as low-dose IL-2, administration of regulatory T cells, or leukemia/virus-specific dendritic cells and/or other specific T cells (9).

## Conclusions

If our hypothesis is confirmed by larger-scale studies, then V-gene reconstitution, and especially the distribution of rare V genes, can serve as a biomarker for the thymic-independent regeneration pathway. As V-gene segment usage profiling is a relatively simple procedure that is widely used and can be performed with limited sampling depth (20,41,42), we propose that to test the evolution of the V gene usage in the recipients can serve as a simple functional measure of immune recovery that may guide clinicians in monitoring immune recovery post allo-BMT transplant.

## Methods

We compared multiple T-cell repertoires as a means to follow T cell repertoire changes post allo-BMT in HLA-matched related donor and recipient pairs. We first characterize each repertoire by either its CDR3 beta or V-gene distribution and then compared these characteristics in all other repertoires (See flowchart of analyses in Figure 1). We used this method to study the changes in post allo-BMT TCRβ repertoire and show how V gene usage provides information on the similarities and differences between the donor and the recipient’s developing immune systems.

## Subjects and samples

The data used for this study was derived from a prospective clinical study (NCT00809276) by Kanakry et al (1). The dataset is freely available at the Adaptive Biotechnologies database (Seattle, WA; https://clients.adaptivebiotech.com).

TCR repertoires from four pairs of individuals were analyzed; each pair is composed of a donor and a recipient of bone marrow from the same family. In addition, seven patients did not have pre-transplant sequencing data for all time points (TP) post-transplant, so only repertoires from one month and one-year post-transplant were compared. Donor-recipient pair details are available in Table 1.

All recipients were treated with myeloablative treatment of 40 mg/m^2^/day IV Fludarabine plus Busulfan based on real-time PKs to achieve a target concentration of 900 ng/mg. IV daily for 4 days prior to transplantation and high-dose post-transplantation cyclophosphamide (PTCy), which effectively reduces graft-versus-host disease, on days 3 and 4 post-transplantation as single-agent for GVHD prophylaxis (43). Donors provided peripheral blood samples on the day of bone marrow harvesting (1). The analyses focused on TCR β chain sequencing data of peripheral blood mononuclear cells (PBMC) samples taken from the donors and from recipients at time points 0 (transplantation),1, 12 and 36-48 months post-transplantation.

CDR3 beta amino acid sequences and V-gene segments were extracted from the sequencing files after removing out-of-frame sequences. In the V-gene frequency analysis, we ignored all sequences with ‘unresolved’ V-gene. We did not analyze J genes as they are rather uniform and thus supply very limited information.

We used the processed data by the authors in the original manuscript(1). Similar clones were merged in the original analysis. In the processed data there are practically no clones with equal CDR3 sequences, as can seen by the distance distribution between clones in samples from the different time points (Figure S1). The total number of clones in each sample is shown in Figure S2.

We randomly picked a sample of 1000 clones per file. One file (037, 1-month post-transplant) does not contain enough data (it contains 767 sequences) so we used it as is.

We here propose a relatively simple means of comparing multiple T-cell repertoires. We utilize a 1,000 randomly picked sequences from each repertoire. While T-cell repertoires are theoretically huge, in practice they have an order of 1.e6-1.e7 clones. Moreover, our sampled repertoires represent mainly the larger clones, which are typically between 1,000 and 10,000 clones in the vast majority of samples. Indeed, we used 1,000 clones that constitute a big fraction of the total number of clones (See Figure S2 for the total number of clones in each sample).

### CDR3 sequence distance

We analyzed two aspects of the TCR repertoire: CDR3 beta sequences and V-gene distribution. Analysis flow chart is illustrated in Figure 1.

To find the level of similarity (referred to here as ‘distance’) in CDR3 beta composition between samples, we calculated for each CDR3 sequence in every sample its distance from all sampled CDR3 sequences in another sample and found its nearest neighbor (i.e. the sequence in the other sample with the lowest edit distance). We then computed the average edit distance of these closest neighbors. The Edit Distance used here was the Levenshtein distance (44). The Levenshtein distance represents the number of insertions, deletions, and substitutions required to change one word to another (45). When computing the self-distance, which is the distance between S and S, the diagonal values (edit distances between the same sequences – all zeros) were ignored.

### V gene usage

The V-gene distributions of each sample (shown in Figure S3) were first found by a previously used methodology (46,47). Generally, some V genes are highly used, such as V02-01,V05-01 and V19-01, while others are very rarely used (48). These very frequent V genes are often not distinctive as they bias the computation of the absolute distance between V gene usages and often mask the signal distinguishing between individuals and time points in T cell reconstruction. To avoid this bias, we employed a symmetrized version of the Kullback-Leibler (KL) divergence (D(p,q)+D(q,p))(35) (equation 1). The KL divergence measures the difference between two probability distributions, where KL of 0 indicates that the two distributions are identical. To avoid zero occurrences of V-segments, we have used a pseudo-count of 1.e-4. Although the pseudo count affects the KL, it has a very limited effect on the difference between distances (See figure S4 in supplementary for multiple random comparisons between pairs of samples using different constants).

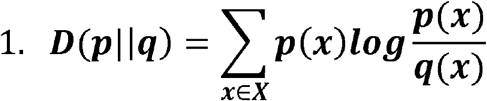

Where p and q are the two probability distributions and x is one value out of a set of possibilities X. We also measured the distance between two V distributions with Euclidian distance (Equation 2).

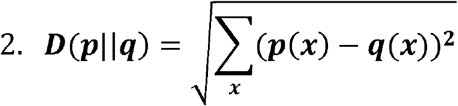

### MDS and clustering

For visualization, a multidimensional scaling (classical MDS) algorithm was used; MDS is a form of non-linear dimensionality reduction used to display distances. In our case, the distances were between each pair of CDR3 sequences (derived from the calculation of the ED), or the distances between each two V genes probability distributions (derived from the calculation of the KL given in equation 1).

MDS places samples in a multi-dimensional space such that the between-sample distances are preserved as well as possible (49). The results here are presented with a three dimensional MDS.

Clustering of samples was performed with Hierarchical cluster analysis (HCA) (50) on the KL and ED distances matrixes (‘scipy.cluster.hierarchy.linkage’ and ‘scipy.cluster.hierarchy.dendogram’ packages).

### Code

All analyses were performed in Python language, version 3.6. The edit distance between two sequences was calculated using ‘jellyfish.levenshtein_distance’ package, Euclidean distance between two V distributions was calculated with package ‘numpy.linalg.norm’ and MDS was calculated with ‘sklearn.manifold.MDS’ package.

### Statistical analysis

Statistical analyses were performed in Python language, version 3.6. The difference between the four groups in our analysis (detailed below) was tested using ANOVA (‘scipy.stats.f_oneway’ package). If the ANOVA was significant, we performed paired Student’s t-tests to compare the two groups d (‘scipy.stats.ttest_ind’ package). A Benjamini Hochberg False Discovery Rate correction(51) was performed for multiple comparisons with three comparisons in figure 2C and six comparisons in 3A.

We compared four groups of samples pairs:

- The baseline group composed of all recipients (R) immediately post-transplant. All samples are at the same time point (TP). Intra-sample distance is 0 here per definition but is required as baseline for the following analyses. The group is referred to as (TP+R+).
- A group composed of pairs of different recipients whose repertoires are compared at the same time point (TP+R-).
- A group composed of the repertoires from the same recipient at different time points (TP-R+).
- A group composed of pairs of different recipients whose repertoires are compared at different time points (TP-R-).

An ANOVA test was performed with all groups and also after excluding the first group of ‘same time point, same recipient’ due to the low mean. Then, Student’s t-tests were performed between all group pairs in order to find significantly different groups.

#### Regression model

The relationship between the edit distance and its components can be described by a positive regression problem:

Distance (recipient1 at TP1, recipient2 at TP2) = Distance (recipient1, recipient2) + Distance (TP1, TP2) + noise (Equation 3).

To understand how the edit distance between the samples (left side of the equation) is influenced by the distance between individuals and by the time point (right side of the equation), we performed non-negative regression analysis with Python package ‘scipy.optimize.nnls’ which computes a non-negative least squares.

## Supporting information

All supplementary figures

## Availability of data and materials

The datasets generated and/or analysed during the current study are available in the Adaptive Biotechnologies database (Seattle, WA; https://clients.adaptivebiotech.com).

## Author Contributions

1. SD designed the formalism and implemented it.
2. NTM collected the data and wrote the manuscript.
3. AR and AAK collected the data and prepared it.
4. SE supervised the work.
5. YL supervised the work and wrote the manuscript.

## References

1. Kanakry CG, Coffey DG, Towlerton AMH, Vulic A, Storer BE, Chou J, et al. Origin and evolution of the T cell repertoire after posttransplantation cyclophosphamide. JCI Insight. 2016;1(5):1–33.

2. Yew PY, Alachkar H, Yamaguchi R, Kiyotani K, Fang H, Yap KL, et al. Quantitative characterization of T-cell repertoire in allogeneic hematopoietic stem cell transplant recipients. Bone Marrow Transplant. 2015;50(9):1227–34.

3. Robinson TM, O’Donnell P V, Fuchs EJ, Luznik L. Haploidentical bone marrow and stem cell transplantation: experience with post-transplantation cyclophosphamide. Semin Hematol. 2016 Apr;53(2):90–7.

4. Gauthier S, Moutuou MM, Daudelin F, Leboeuf D, Guimond M. IL-7 Is the Limiting Homeostatic Factor that Constrains Homeostatic Proliferation of CD8 + T Cells after Allogeneic Stem Cell Transplantation and Graft-versus-Host Disease. Biol Blood Marrow Transplant. 2018;000:1–8.

5. Fallen PR, Mcgreavey L, Madrigal JA, Potter M, Ethell M, Prentice HG, et al. Lymphoid Reconstitution Factors affecting reconstitution of the T cell compartment in allogeneic haematopoietic cell transplant recipients. Bone Marrow Transplant. 2003;32(10):1001–14.

6. Sakai R, Kinouchi T, Kawamoto S, Dana MR, Hamamoto T, Tsuru T, et al. Construction of human corneal endothelial cDNA library and identification of novel active genes. Invest Ophthalmol Vis Sci. 2002;43(6):1749–56.

7. Lynch HE, Goldberg GL, Chidgey A, Brink MRM Van Den, Boyd R, Sempowski GD. Thymic involution and immune reconstitution. Cell. 2009;30(7):366–73.

8. Toubert A, Glauzy S, Douay C, Clave E. Thymus and immune reconstitution after allogeneic hematopoietic stem cell transplantation in humans□: never say. Tissue Antigens. 2012;79(2):83–9.

9. Koning C De, Nierkens S, Boelens JJ. Strategies before, during, and after hematopoietic cell transplantation to improve T-cell immune reconstitution. Blood. 2016;128(23):2607–16.

10. Chaudhry MS, Velardi E, Malard F, Brink MRM Van Den. Immune Reconstitution after Allogeneic Hematopoietic Stem Cell Transplantation: Time To T Up the Thymus. J Immunol. 2019;198:40–6.

11. Krenger W, Blazar BR, Holla GA. Thymic T-cell development in allogeneic stem cell transplantation. Blood. 2011;117(25):6768–77.

12. Bassing CH, Swat W, Alt FW. The Mechanism and Regulation of Chromosomal V (D) J Recombination. Cell. 2002;109(D):45–55.

13. Schrama D, Ritter C, Becker JC. T cell receptor repertoire usage in cancer as a surrogate marker for immune responses. Semin Immunopathol. 2017;39:255–68.

14. Benichou JIC, van Heijst JWJ, Glanville J LY. Converging evolution leads to near maximal junction diversity through parallel mechanisms in B and T cell receptors. Phys Biol. 2017;14(4):045003.

15. Lissina A, Chakrabarti LA, Takiguchi M, Appay V. TCR clonotypes: molecular determinants of T-cell efficacy against HIV. Curr Opin Virol. 2016;16:77–85.

16. Cells HCDT, Luo W, Ma L, Wen Q, Zhou M, Wang X. Analysis of the Conservation of T Cell Receptor Alpha and Beta Chain Variable Regions Gene in pp65 Peptide-Specific. 2009;(2):105–10.

17. Shugay M, Bolotin DA, Putintseva EV, Pogorelyy MV, Mamedov IZ CD. Huge overlap of individual TCR beta repertoires. Front Immunol. 2013;4(DEC):1–3.

18. Britanova OV, Putintseva EV, Shugay M, Merzlyak EM, Turchaninova MA, Staroverov DB, Bolotin DA, Lukyanov S, Bogdanova EA, Mamedov IZ, Lebedev YB CD. Age-Related Decrease in TCR Repertoire Diversity Measured with Deep and Normalized Sequence Profiling. J Immunol. 2014;192(6):2689–98.

19. Burrows SR, Silins SL, Moss DJ, Khanna R, Misko IS AV. T Cell Receptor Repertoire for a Viral Epitope in Humans Is Diversified by Tolerance to a Background Major Histocompatibility Complex Antigen. J Exp Med. 1995;182(6):1703–1715.

20. Madi A, Shifrut E, Reich-Zeliger S, Gal H, Best K, Ndifon W, et al. T-cell receptor repertoires share a restricted set of public and abundant CDR3 sequences that are associated with self-related immunity. Genome Res. 2014;24(10):1603–12.

21. Dean J, Emerson RO, Vignali M, Sherwood AM, Rieder MJ, Carlson CS, et al. Annotation of pseudogenic gene segments by massively parallel sequencing of rearranged lymphocyte receptor loci. Genome Med [Internet]. 2015;123(7):1–8. Available from: http://dx.doi.org/10.1186/s13073-015-0238-z

22. Heather JM, Ismail M, Oakes T, Chain B. High-throughput sequencing of the T-cell receptor repertoire□: pitfalls and opportunities. Brief Bioinform. 2018;19(4):554–65.

23. Benichou J, Ben-Hamo R, Louzoun Y, Efroni S. Rep-Seq□: uncovering the immunological repertoire through next-generation sequencing. Immunology. 2011;135(3):183–91.

24. Muraro PA, Robins H, Malhotra S, Howell M, Phippard D, Desmarais C, et al. Brief report T cell repertoire following autologous stem cell transplantation for multiple sclerosis. 2014;124(3):1168–72.

25. Ogonek J, Kralj Juric M, Ghimire S, Varanasi PR, Holler E, Greinix H WE. Immune Reconstitution after Allogeneic Hematopoietic Stem Cell Transplantation. Front Immunol. 2016;17(7):507.

26. Pogorelyy MV, Fedorova AD, McLaren JE, Ladell K, Bagaev DV, Eliseev AV, Mikelov AI, Koneva AE, Zvyagin IV, Price DA, Chudakov DM SM. Exploring the pre-immune landscape of antigen-specific T Cells. Genome Med. 2018;10(1):68–82.

27. Gerlinger M, Quezada SA, Peggs KS, Furness AJ, Fisher R, Marafioti T, et al. Ultra-deep T cell receptor sequencing reveals the complexity and intratumour heterogeneity of T cell clones in renal cell carcinomas. J Pathol. 2013;231(4):424–32.

28. Cieri N, Oliveira G, Greco R, Forcato M, Taccioli C, Cianciotti B, et al. Generation of human memory stem T cells after haploidentical T-replete hematopoietic stem cell transplantation. Blood. 2015 Apr;125(18):2865–74.

29. Mamedov IZ, Britanova OV, Bolotin DA, Chkalina AV, Staroverov DB, Zvyagin IV, Kotlobay AA, Turchaninova MA, Fedorenko DA, Novik AA, Sharonov GV, Lukyanov S, Chudakov DM LY. Quantitative tracking of T cell clones after haematopoietic stem cell transplantation. EMBO Mol Med. 2011;3(4):201–7.

30. Nakasone H, Tanaka Y, Yamazaki R, Terasako K, Sato M, Sakamoto K, et al. Single-cell T-cell receptor-β analysis of HLA-A*2402-restricted CMV-pp65-specific cytotoxic T-cells in allogeneic hematopoietic SCT. Bone Marrow Transplant. 2014;49(1):87–94.

31. Izraelson M, Nakonechnaya TO., Moltedo B, Erogov ES., Kasatskaya SA, Ekaterina V, Mamedov IZ, Staroverov DB, Irina I, et al. Comparative analysis of murine T-cell receptor repertoires. Immunology. 2018;153(2):133–44.

32. Elhanati Y, Sethna Z, Callan CG, Mora T, Walczak AM. Predicting the spectrum of TCR repertoire sharing with a data-driven model of recombination. Immunol Rev. 2018;284(1):167–79.

33. Meier JA, Haque M, Fawaz M, Abdeen H, Coffey D, Towlerton A, et al. Biology of Blood and T Cell Repertoire Evolution after Allogeneic Bone Marrow Transplantation□: An Organizational Perspective. Biol Blood Marrow Transplant [Internet]. 2019;000:1–15. Available from: https://doi.org/10.1016/j.bbmt.2019.01.021

34. Turner SJ, Kedzierska K, La Gruta NL, Webby R, Doherty PC. Characterization of CD8+ T cell repertoire diversity and persistence in the influenza a virus model of localized, transient infection. Semin Immunol. 2004;16(3):179–84.

35. Kullback, S.; Leibler RA. On Information and Sufficiency. Ann Math Stat. 1951;22(1):79–86.

36. Koning D, Costa a. I, Hoof I, Miles JJ, Nanlohy NM, Ladell K, et al. CD8+ TCR Repertoire Formation Is Guided Primarily by the Peptide Component of the Antigenic Complex. J Immunol. 2012;190(3):931–9.

37. Douek DC, Vescio RA, Betts MR, Brenchley JM, Hill BJ, Zhang L, et al. Early report Assessment of thymic output in adults after haematopoietic stem-cell transplantation and prediction of T-cell reconstitution. Lancet. 2000;355:1875–81.

38. Ringhoffer S, Rojewski M, Döhner H, Bunjes D, Ringhoffer M. T-cell reconstitution after allogeneic stem cell transplantation□: assessment by measurement of the sjTREC / β TREC ratio and thymic naïve T cells. Haematologica. 2013;98(10):1600–8.

39. Dickinson AM. Immune Reconstitution after Allogeneic Hematopoietic Stem Cell Transplantation. Front Immunol. 2016;17(7):507.

40. Buhler S, Bettens F, Dantin C, Marc SF, Mamez AA, Yves SM, et al. Genetic T-cell receptor diversity at 1 year following allogeneic hematopoietic stem cell transplantation. Leukemia. 2019;34(5):1422–32.

41. Nakano N. T cell receptor V gene usage of islet beta cell-reactive T cells is not restricted in non-obese diabetic mice. J Exp Med. 1991 May 1;173(5):1091–7.

42. Scheckelhoff MR, Deepe GS. Pulmonary V beta 4+ T cells from Histoplasma capsulatum-infected mice respond to a homologue of Sec31 that confers a protective response. J Infect Dis. 2006;193(6):888–97.

43. Kanakry CG, O’Donnell P V., Furlong T, de Lima MJ, Wei W, Medeot M, et al. Multi-Institutional Study of Post-Transplantation Cyclophosphamide As Single-Agent Graft-Versus-Host Disease Prophylaxis After Allogeneic Bone Marrow Transplantation Using Myeloablative Busulfan and Fludarabine Conditioning. J Clin Oncol. 2014 Nov;32(31):3497–505.

44. Levenshtein VI. Binary codes capable of correcting deletions, insertions, and reversals. Sov Phys Dokl. 1966;10(8):707–710.

45. Navarro Gonzalo. A Guided Tour to Approximate String Matching. ACM Comput Surv. 2001;33(1):31–88.

46. Zvyagin IV, Pogorelyy MV, Ivanova ME, Komech EA, Shugay M, Bolotin DA, Shelenkov AA, Kurnosov AA, Staroverov DB, Chudakov DM, Lebedev YB MI. Distinctive properties of identical twins’ TCR repertoires revealed by high-throughput sequencing. Proc Natl Acad Sci. 2014;111(16):5980–5.

47. Rubelt F, Bolen CR, Mcguire HM, Heiden JA Vander, Gadala-maria D, Levin M, et al. Individual heritable differences result in unique cell lymphocyte receptor repertoires of na&iuml;ve and antigen-experienced cells. Nat Commun [Internet]. 2016;6:1–12. Available from: http://dx.doi.org/10.1038/ncomms11112

48. Robins H. Overlap and effective size of the human CD8+ T-cell receptor repertoire. Sci Transl Med. 2010;2(47):1–9.

49. Borg, I., Groenen P. Modern Multidimensional Scaling: theory and applications. New York Springer-Verlag. 2005;2nd Editio:207–212.

50. Johnson SC. Hierarchical clustering schemes. Psychometrika. 1967;32:241–254.

51. Benjamini Y, Drai D, Elmer G, Kafkafi N GI. Controlling the false discovery rate in behavior genetics research. Behav Brain Res. 2001;125(1–2):279–84.

