## Supplementary material for "CDR3 and V- genes show distinct reconstitution patterns in T-cell repertoire post allogeneic bone marrow transplantation": All supplementary figures

### Supplementary Figure S1

Distance distribution between clones in samples from the different time points. It shows that in the processed data there are practically no clones with equal CDR3 sequences in different samples. The distance definition employs Gonnet et al.'s measures of multiple sequence alignment(1).

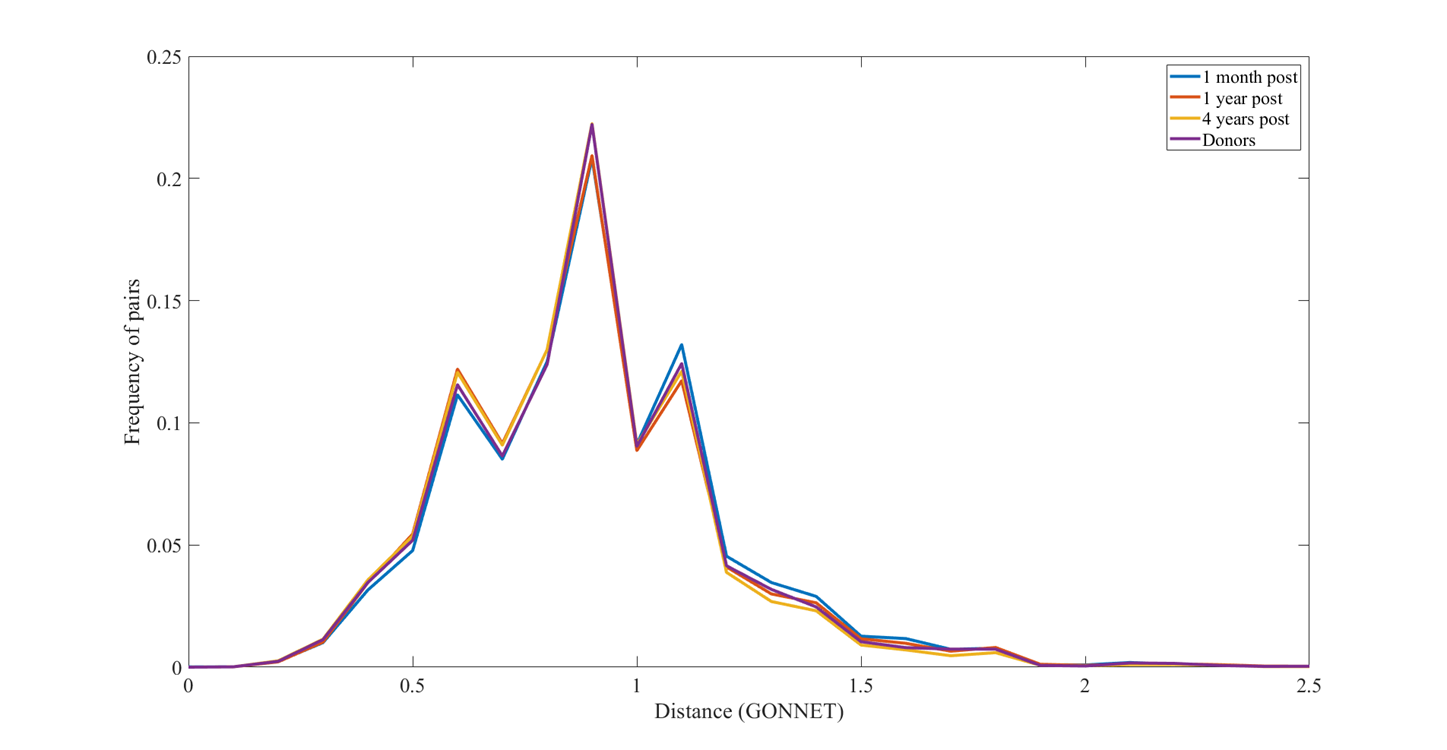

**Supplementary figure S2**

The total number of clones in each sample

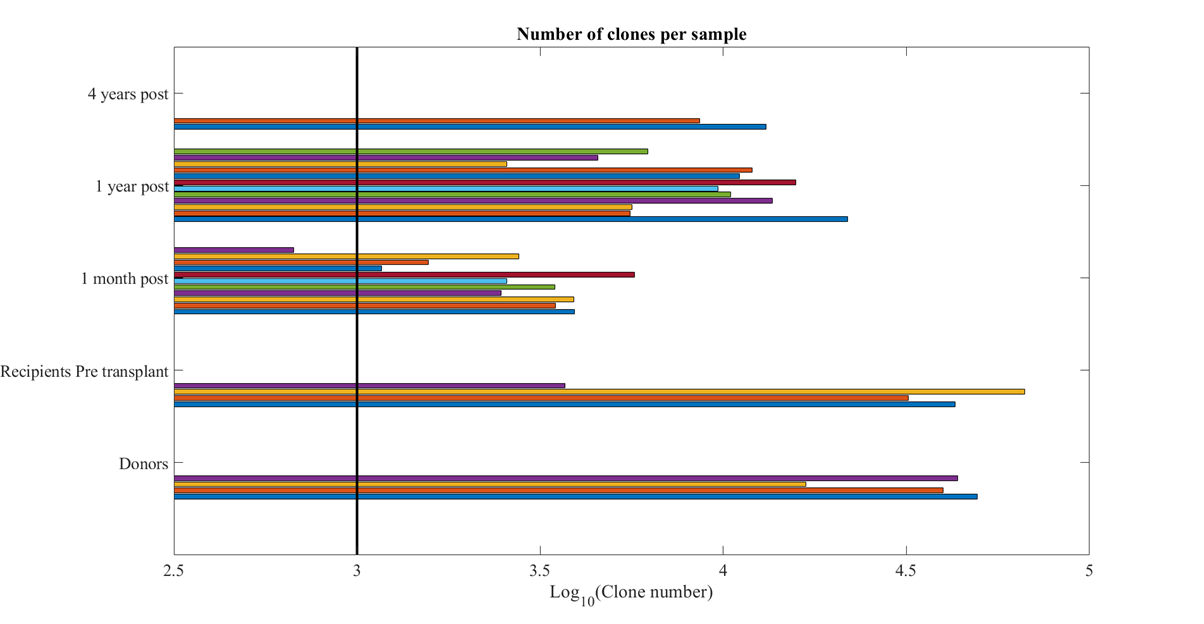

### Supplementary figure S3

### V-gene usage: V-gene frequencies over all samples. Each color stands for a different sample (each sample was taken from an individual recipient at a single time point).

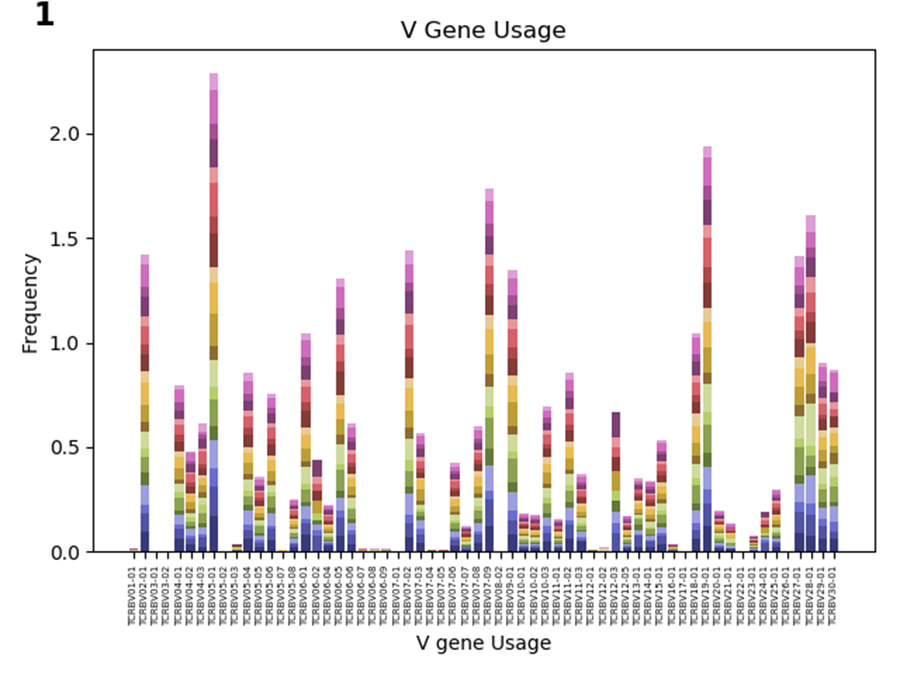

### Supplementary figure S4

Although the pseudo count affects the KL, it has a very limited effect on the difference between distances, as can be seen by multiple random comparisons between pairs of samples using different constants. Each line represents a different pair of distances. The x axis represents the constant added to the KL divergence. The samples compared are the 1 month and 1 year post transplant for the first patient, 4 year post transplant of patient 2 and donor 2.This choice of samples is arbitrary and other sample pairs give similar results.

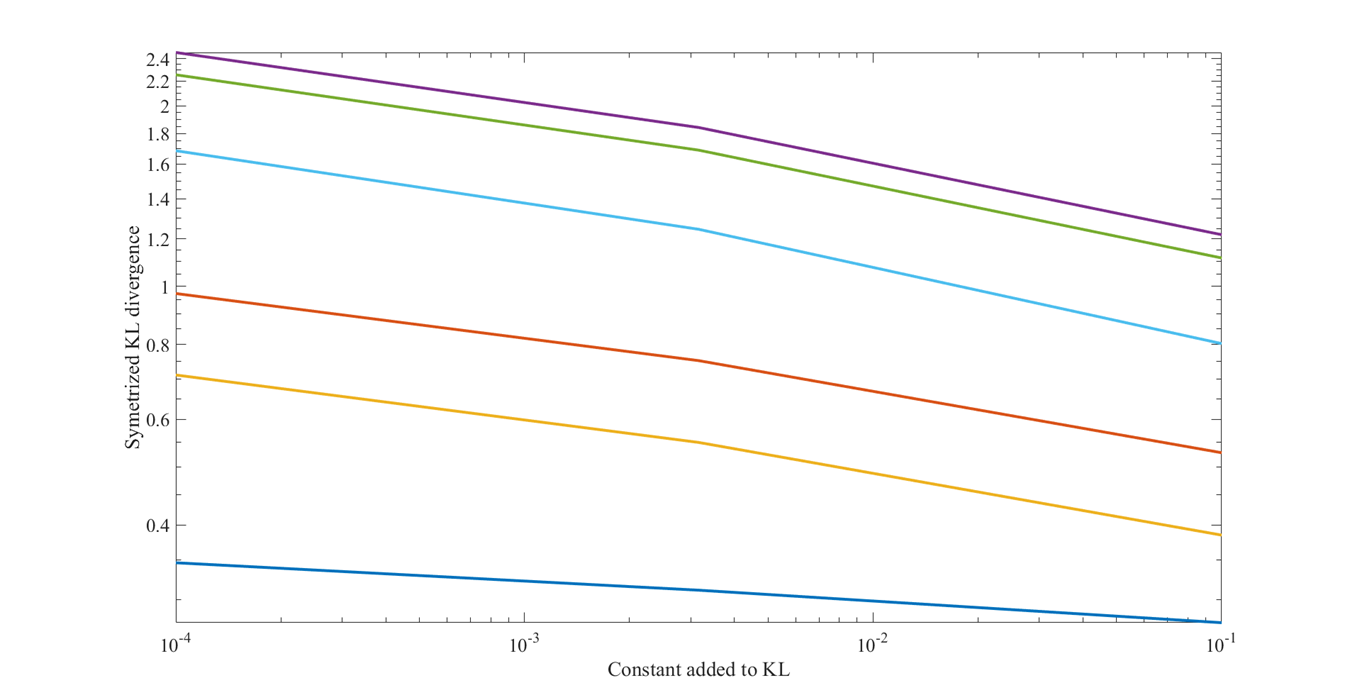

### Supplementary figure S5

Beyond a sampling of 1,000 clones (x axis in figure below), the V gene usage is conserved, as seen by the unchanged colors to the right of the black line. Each row is a different V-gene. Not all V-gene names are marked for the sake of figure clarity.

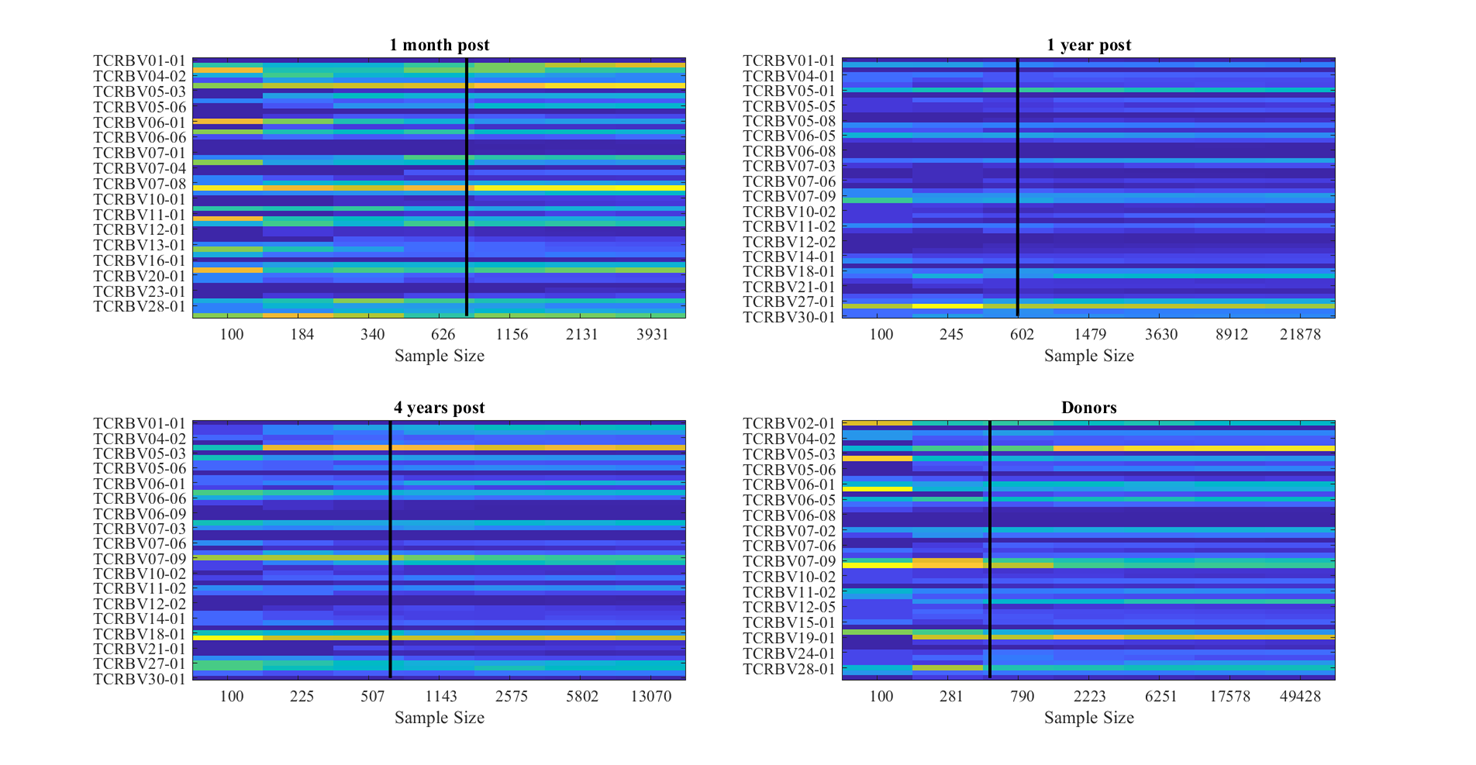

**Supplementary Figure S6**

We plotted the results for each patient to show that they are highly consistent in their pattern of decreasing distance to pre-transplant recipient, and increasing distance to donor.

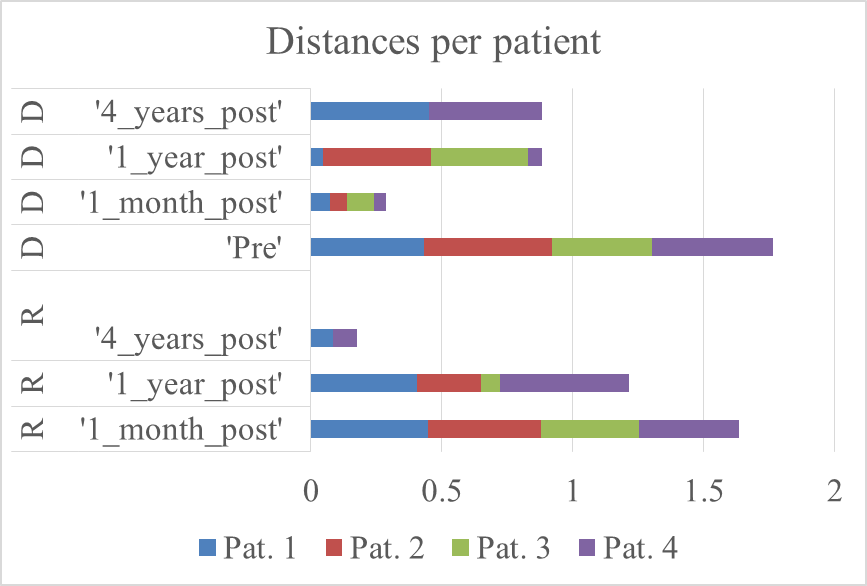

### 4 Supplementary Figure S7

### A. V-gene Hierarchical clustering. The Euclidian distances between samples were clustered with 'single' method. The first half of the labels indicates one of the five states: Donors (D), Pre-transplant (P), 1-month post (1M), 1-year post (1Y) and 3-4 years post-transplant (4Y). The second half of the labels indicates the ID of the recipient. Labels are colored according to time points.

### B. In the Euclidian V-gene distribution heatmap each cube represents the Euclidian distance from two distributions of V-gene usage. Blue indicates smaller values, i.e., closer samples. Samples are positioned by the order of the dendrogram leaves from 1A.

### C. A swarm plot (i.e., a categorical scatterplot) of Euclidian distances from 1B, grouped into 4 categories: 'same time point (TP), same recipient' (Blue), 'same TP, different recipient' (Green), 'different TP, same recipient' (Red) and 'different TP, different recipient' (Purple). The t-test's p-value is indicated as ** (p-value < 0.01) and *** (p-value < 0.001).

### D. MDS of the Euclidian distance matrix from 1B; Different time points are shown in different colors and different IDs are indicated with a variety of shapes.

### E. Euclidian distances of four recipients (011, 037, 016, 019) from their pre-transplant and donors repertoires as a function of time.

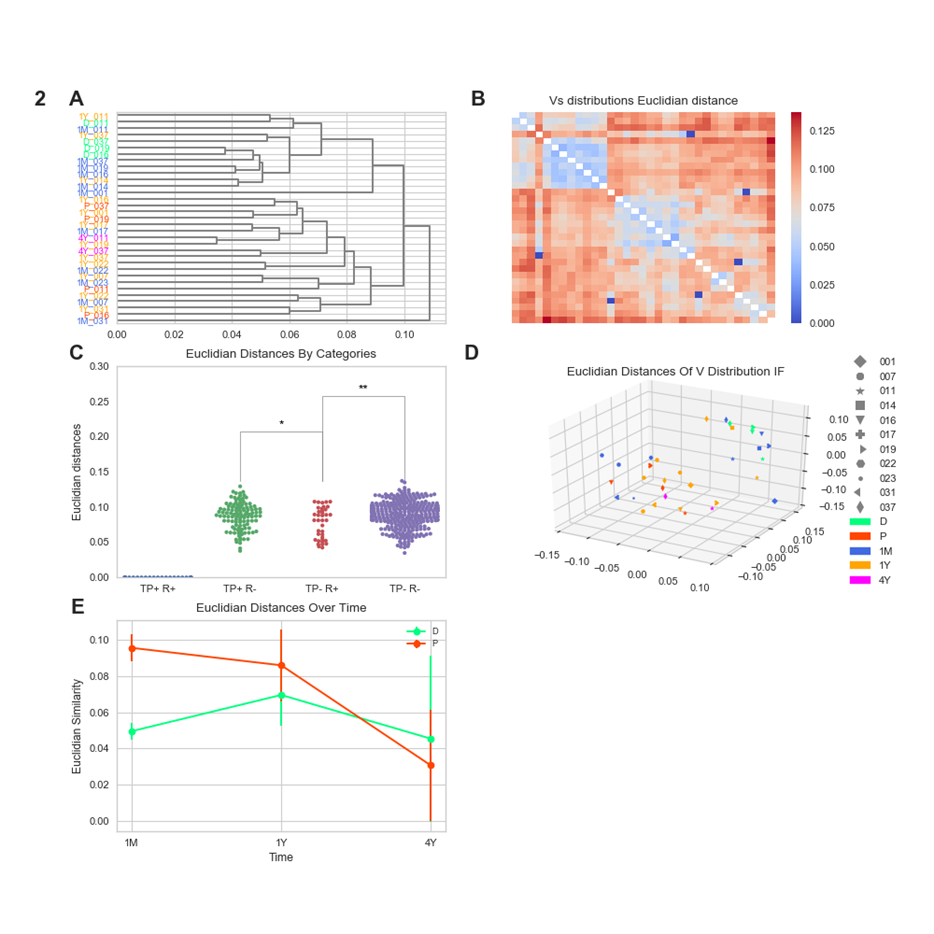

**Reference**

1.	Gonnet GH, Korostensky C BS. Evaluation measures of multiple sequence alignments. J Comput Biol. 2000;7(1–2):261–76.
